# *In Vitro* Generation and Characterization of The Wu Syndrome Model That Causes Mental Retardation in Neural Cell Lines

**DOI:** 10.1101/2024.07.01.601494

**Authors:** Sumeyye Seher Karaman, Tuba Sevik, Selin Akdemir, Burak Bilgin, Esra Karadeli, Dilay Yalcin, Buse Baran, Beste Gelsin, Buket Budaklar, Berranur Sert, Gamze Gulden, Cihan Tastan

## Abstract

Wu Syndrome, also known as X-Linked Wu Type Intellectual Developmental Disorder, is caused by a mutation in the GRIA3 (Glutamate Ionotropic Receptor AMPA Type Subunit 3) gene located at position 25 on the X chromosome. GRIA3 encodes iGluR3, a subunit of the AMPA (α-amino-3-hydroxy-5-methyl-4-isoxazolepropionic acid) receptor, which plays a critical role in rapid excitatory synaptic transmission in the central nervous system. This receptor is essential for learning, memory, and the processes of long-term depression (LTD) and long-term potentiation (LTP). Despite its significance, Wu Syndrome remains under-researched and lacks effective treatments. Notably, some genetic variants have been identified, but many, including the W637S variant, are still unstudied. This study pioneers the development of a Wu Syndrome model in neural cell lines using genetic modification techniques to identify and characterize new GRIA3 variants. By focusing on variants such as G833R and W637S, this research provides novel insights into their effects on GRIA3 function, paving the way for potential therapeutic strategies. This is the first study to explore the responses of neural cells to these mutations in vitro, thereby contributing valuable knowledge toward understanding and treating Wu Syndrome.

## INTRODUCTION

Wu Syndrome, an X-Linked Wu Type Intellectual Developmental Disorder, is caused by a mutation in the GRIA3 (Glutamate ionotropic receptor AMPA type subunit 3) gene located at location 25 on the X chromosome. GRIA3 encodes iGluR3 (Ionotropic glutamate receptor 3), one of the subunits of the AMPA (α-amino-3-hydroxy-5-methyl-4-isoxazolepropionic acid) receptor. In the central nervous system, AMPA receptors function as the primary mediators of rapid excitatory synaptic transmission, critical for learning, and memory, as well as the initiation and continuation of LTD (long-term depression) and LTP (long-term potentiation). Wu Syndrome is a rare disease that has not been adequately studied in the literature. There is still no treatment, and although some variants have been studied, others are yet to be discovered. The effects of these unstudied variants on the function and characterization of GRIA3 are unknown. Wu Syndrome is among genetic disorders that are challenging to study due to its rarity. Although it has been treated with antiepileptic drugs, these interventions have only temporarily prevented seizures (Sun et al., 2021).

While the abilities of thinking, reasoning, perceiving objective facts, comprehending, judging, and drawing conclusions are called intelligence, retardation is observed in adapting to daily life and cognitive functions in mental retardation. It begins before the age of 18 and its effects last a lifetime. An IQ (intelligence quotient) below 70 has been determined. It is a disability characterized by a marked limitation of conceptual, social, and adaptive skills (Karadag et al., 2017). Intellectual disability (ID) can be caused by environmental factors as well as genetic reasons. Wu Syndrome develops due to the X chromosome and it is transferred as X-linked Recessive Inheritance. Defects in the X-chromosome and thus an excess of males have been reported. However, although rare, it has also been reported in female patients (Philips et al., 2014).

GRIA3 is the gene encoding iGLUR3, one of the four AMPA receptors. It is located at Xq25, band 25 on the long arm of the X chromosome. The GRIA3 gene has 16 exons. In humans, mutations in the GRIA3 gene cause X-linked mental retardation. Because it is a mutation on the X chromosome, pathogenic variants are mostly seen in male patients, although cases in females have also been rarely reported. Mutations in the GRIA3 gene lead to the loss or gain of iGluR3 function and channel current. As a result, abnormal memory formation and learning difficulties occur. Behavioral disorders, wakefulness, and sleep attacks are also observed. The brain stem, medulla, olfactory bulb, amygdala, cerebellum, striatum, hippocampus, cerebral cortex, and thalamus are locations of AMPA receptors (AMPAR) that include iGluR3. These regions are also involved in epileptic activity, so iGluR3 dysfunction may also manifest epileptic features (Martinez et al., 2022; Pei et al., 2009).

Glutamate-gated ion channels called AMPA receptors facilitate fast excitatory synaptic transmission in the CNS. It has been reported that they participate in synaptic plasticity, an essential process for brain function. There are four genes encoding the four types of AMPARs: GRIA1, GRIA2, GRIA3, and GRIA4. AMPA receptors are composed of four subunits, each with a homomeric or heteromeric combination of iGluR1, iGluR2, iGluR3, and iGluR4. Once activated by the binding of glutamate, AMPA receptors exhibit sodium permeability. Depolarization of the cell is achieved by sodium passage through AMPAR, thus removing the magnesium block of the NMDA receptor (NMDAR) and truly activating NMDAR. For this reason, the AMPA receptor is of critical importance for the activation of the NMDA receptor.

All AMPAR subunits are available in two forms with alternative splicing known as FLIP and FLOP. FLIP and FLOP exhibit a short-range amino acid difference. FLOP variations are expressed after birth, whereas FLIP variants are mostly expressed during pregnancy. In adulthood, their expression levels are nearly equal. These differences have an impact on the susceptibility of heteromeric AMPA receptors to allosteric modulators, as well as the rate and degree of desensitization. The FLIP isoform shows slower desensitization than the FLOP isoform, and the two exhibit different desensitization kinetics. Although the iGluR3 FLOP variant opens the channel at the same rate as the FLIP isoform, it closes the channel four times faster. These variants are assumed to work to adjust the gating balance. In both FLIP and FLOP isoforms, a minimum of two glutamate molecules is required to bind to the channel and cause it to open. The expression of these isoforms varies according to tissue, cell type, and age. FLIP variants predominate before birth, while FLOP variants are expressed after birth. In adulthood, they are almost equal. In some neurological disorders, abnormalities can be observed in the expression of isoforms relative to each other (Vandenberghe et al., 2005; Henley et al., 2022; Lüscher et al., 2012; Kato et al., 2010; Lau et al., 2010; Kew et al., 2005; Alt et al., 2006; Pei et al., 2007).

Given the significant role of AMPA receptors in synaptic transmission and their implications in neurological disorders, it is crucial to deepen our understanding of the various GRIA3 mutations and their functional impacts. This study aims to address the existing knowledge gap by developing a robust in vitro model to investigate the effects of specific GRIA3 variants, particularly G833R and W637S, in neural cell lines. By employing advanced genetic modification techniques, we will explore the cellular responses and pathological mechanisms associated with these mutations. This research not only seeks to enhance our comprehension of Wu Syndrome but also aims to lay the groundwork for future therapeutic interventions. Through detailed characterization of these variants, we hope to contribute to the broader effort of finding effective treatments for this rare and debilitating disorder.

## MATERIALS AND METHODS

### Plasmids and Synthesized Sequences

All plasmids used in this study and selected sequences are listed in **Table 1**. The desired sequences, including the FLIP and FLOP isomers, as well as the G833R and W637S variants of the GRIA3 gene, were ordered from GenScript. Subsequently, these sequences were synthesized by GenScript and cloned into pHIV-EGFP (Addgene, #21373).

**Table 1:**
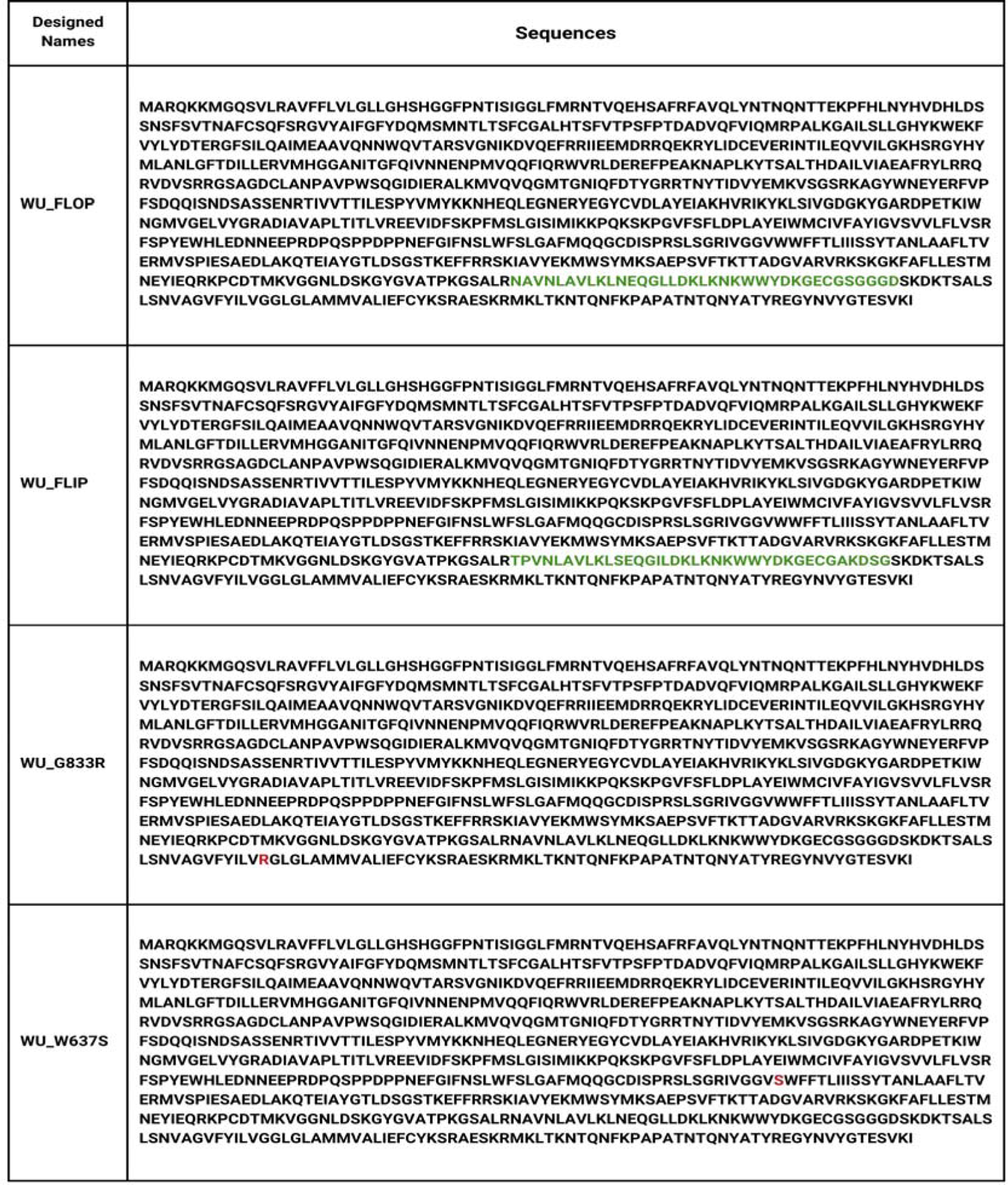
Genetic sequences incorporated into the study. The green color indicates the distinguishing features that separate the isoforms from each other. The change that caused the variation is indicated in red.

### Transformation

Using the heat shock approach, plasmid DNAs are transformed into bacteria that belong to the competitive *Escherichia coli* DH5α strain. On LB agar plates with ampicillin, bacteria are added and then incubated for 16 hours at 37 °C. Selected colonies will be transferred to liquid culture and then plasmid isolation will be performed with the ZymoPURE Plasmid Maxiprep Kit (#D4202, ZYMO).

### Agarose Gel Electrophoresis

To control the isolated plasmids, DNA samples loaded into a 1.5% agarose gel electrophoresis system prepared in 1X TAE (Tris-acetate-EDTA) buffer solution were run at 120V for 45 minutes. The samples were analyzed for supercoiled structures and chromosomal DNA contamination percentages with BIO-RAD Image Lab software.

### Cell Culture

JURKAT cells are cultured in RPMI medium containing 10% fetal bovine serum (FBS), 200 U/ml penicillin/streptomycin antibiotic, 2 mM L-glutamine, 1X MEM vitamin solution, and 1X NEAA (Non-Essential Amino Acids Solution). HEK293T cells are cultured in Gibco DMEM medium with 10% FBS and 1% penicillin/streptomycin, L-Glutamine medium. A172 cells are cultured in DMEM medium with 15% FBS, 200 U/ml penicillin/streptomycin antibiotic, and 2 mM L-glutamine. SH-SY5Y cells are cultured in DMEM medium with 15% FBS, 200 U/ml penicillin/streptomycin antibiotic, 2 mM L-glutamine, and 1% sodium pyruvate. U87 cells are cultured in DMEM medium with 10% fetal bovine serum, 200 U/ml penicillin/streptomycin antibiotic, and 2 mM L-glutamine. All cells are incubated at 37°C with 5% CO2.

### Cell Survival Analysis

Survival analysis was conducted to assess cell viability. The remaining cells were counted and identified using trypan blue (Biological Industries, #03-102-1B). Viability analysis and cell counting were performed using the BIO-RAD TC20 Automated Cell Counter.

### Lentivirus Production

For the production of lentiviruses, plasmid DNA will be separately transfected with pSPAX2 (Addgene #12260) and pVSV-G (Addgene #138479) envelope and packaging plasmid DNA. Lentiviruses will be produced using HEK293T cells by transfecting all plasmids (2:1 pSPAX2:1 pVSV-G) with PEI (Polyethylenimine) transfection reagent (Merten et al., 2016). Supernatants from HEK293T cells will be collected and packaged 72 hours after transfection to produce recombinant lentivirus (Chen et al., 2018; Taştan et al., 2018; Taştan et al., 2020a; Taştan et al., 2020b). Lentiviruses will be concentrated 100x using the LentiX Concentrator (#631232, Takara) to increase virus concentration (Gandara et al., 2018).

### Lentivirus Titration

JURKAT cells were seeded in 96-well U-bottom plates in 100 μL of RPMI. Other wells were set to contain 10 µL, 3 µL, 1 µL, 0.3 µL, 0.1 µL, and 0.03 µL of 100X concentration lentivirus. 50 μL of lentivirus dilutions of each concentration were sequentially transferred to the wells containing Jurkat cells, and the total volume was adjusted to 150 μL. Cells were incubated for 3 days at 37°C, 5% CO2. Flow cytometry (Beckman Coulter, CytoFlex) was used to calculate the titration results.

### Flow Cytometry

All cells were incubated with GLUR3 Antibody (LSBio, Monoclonal Mouse anti-Rat GRIA3 / iGLUR3 Antibody (PE, IF, WB) LS-C654451) to determine iGLUR3 expression levels. Following the collection of cells as 100,000 cells, 1000 μl of staining buffer was added, and the mixture was centrifuged for five minutes at 400 g. After staining with iGLUR3 Antibody, cells were incubated for 30 minutes at 4°C. Then, 1000 μl of staining buffer was added and centrifuged at 400 g for 5 minutes. After the pellet was dissolved using 200 μl of staining buffer, the supernatant was discarded. The cells were then examined using flow cytometry (Beckman Coulter, CytoFlex) to determine the amount of iGLUR3 expression. Fixation/permeabilization procedures were performed using the BDCytofix/Cytoperm Fixation/Permeabilization Kit (Cat. No. 554714). After 100,000 cells were collected, 200 μl of Fixation/Permeabilization solution (BD) was added and incubated at 4°C for 20 minutes. After washing with BD Perm/Wash™ Buffer (BD), cells were stained with GLUR3 Antibody (LSBio, Monoclonal Mouse anti-Rat GRIA3 / iGLUR3 Antibody (PE, IF, WB) LS-C654451) and incubated at 4°C for 30 minutes. Cells were washed with BD Perm/Wash™ Buffer (BD) and analyzed by flow cytometry (Beckman Coulter, CytoFlex) to assess the percentage and the Mean Fluorescence Intensity (MFI) level of GRIA3 expression.

### mRNA Isolation

Cells were harvested and counted to obtain a total of 1×10^6 cells. These cells were then transferred to an RNase-free tube and centrifuged at 2000g for 5 minutes. The supernatant was discarded, and the pellet was resuspended in 0.6 mL of 2-mercaptoethanol-containing lysis buffer. The mixture was vortexed until the pellet was completely dissolved. The homogenized sample was transferred to a new tube and centrifuged at 12000g for 2 minutes. The supernatant was carefully removed and transferred to another RNase-free tube. An equal volume of 1:70 ethanol was added to the supernatant and mixed thoroughly to precipitate the RNA. Next, 700 µL of the total volume was transferred to a spin cartridge and centrifuged at 12000g for 15 seconds. The flow-through was discarded, and the process was repeated until the entire sample had passed through the spin cartridge. Following this, 700 µL of wash buffer I was added to the cartridge, and the mixture was centrifuged again at 12000g for 15 seconds. The flow-through was discarded. Subsequently, 500 µL of wash buffer II was added to the spin cartridge and centrifuged at 12000g for 15 seconds. This washing step was repeated twice to ensure the removal of any contaminants. Finally, the RNA was eluted by adding 30 µL of RNase-free water to the center of the cartridge membrane, incubating for 1 minute, and centrifuging at 12000g for 2 minutes. The eluted RNA was quantified and stored at −20°C until further use.

### cDNA Synthesis

Utilized the Applied Biosystems™ High-Capacity cDNA Reverse Transcription Kit following the specified protocol for a 20 µL reaction volume with up to 2 µg of total RNA. The reaction conditions were 25°C for 10 minutes, 37°C for 120 minutes, 85°C for 5 minutes, followed by a hold at 4°C.

### Reverse Transcriptase Quantitative Real-Time PCR (RT-qPCR)

RT-qPCR was conducted using the initial three primer sets from Bonnet et al. (2009) (Table 2). Each reaction had a final volume of 20 µL. We added 10 µL of Applied Biosystems™ PowerUp™ SYBR™ Green Master Mix to each tube. cDNA amounts were standardized to 100 ng. From the prepared cDNA dilutions, 1 µL was used per reaction. Primer concentrations were maintained between 300-800 nM, with 0.8 µL from a 1:10 primer dilution added to each reaction. Both reverse and forward primers were included in the same tube, and the remaining volume was adjusted with 7.4 µL of sterile water to achieve a total volume of 20 µL per reaction.

**Table 2.**
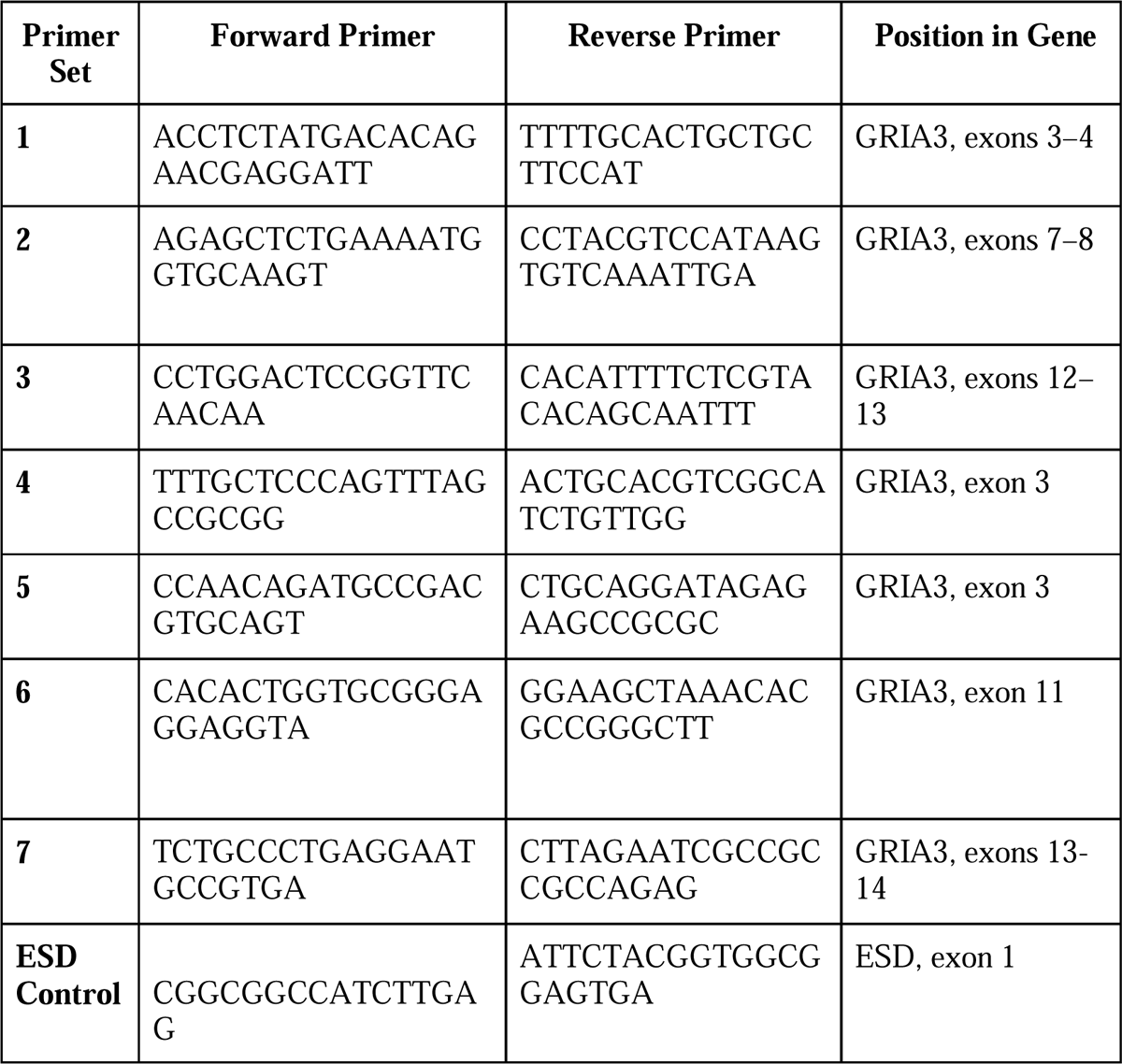
Primer Sets for Amplification of GRIA3 and ESD Control Regions. Primer sets are used for amplifying specific exons of the GRIA3 gene and the ESD control region. Primer sets 4, 5, 6, and 7 are designed to specifically target the FLIP, FLOP isoforms, and W637S, G833R variants of the GRIA3 gene. Primer sets 4, 5, 6, and 7 were designed specifically for the FLIP, FLOP isoforms, and W637S, G833R variants.

The thermal cycling conditions for the non-transduced cells, using primer sets 1, 2, and 3 from Table 2, were as follows: initial denaturation at 95°C for 20 seconds, followed by 40 cycles of 95°C for 3 seconds, 60°C for 30 seconds, and 72°C for 10 seconds. For the transduced cells, primer sets were specifically designed for the FLIP, FLOP, G833R, and W637S variants, corresponding to primer sets 4, 5, 6, and 7 in Table 2. The thermal cycling conditions for these transduced cells were: initial denaturation at 95°C for 20 seconds, followed by 40 cycles of 95°C for 3 seconds, 65°C for 30 seconds, and 72°C for 10 seconds. The ESD gene served as the control amplicon, and all samples were run in duplicate.

### Statistical Approach

CytExpert (Beckman Coulter) was used for the analysis of the flow cytometry data. Statistical analyses were performed using GraphPad Prism 10.1.2 software (GraphPad Software, Inc., San Diego, CA, USA) with a two-tailed t-test using independent mean values. Error bars represent the Standard Error of Means (SEM). For all experiments, significance was defined as p<0.05 and NS= Non-Significant.

## RESULTS

### FLIP-FLOP Isoforms and Variants

Due to the mutually exclusive exon type of alternative splicing, all AMPAR subunits have two versions known as “FLOP” and “FLIP” at exon 14 (**Figure 1a**). FLIP-FLOP isoforms are generated with a variation of 9 amino acids in the extracellular binding domain between the M3 and M4 transmembrane domains, although it contains 38 amino acid sequences in total (**Figure 1b**) (Vandenberghe et al., 2005; Henley et al., 2022; Lüscher et al., 2012; Kato et al., 2010; Lau et al., 2010; Kew et al., 2005; Alt et al., 2006; Pei et al., 2007).

**Figure 1:**
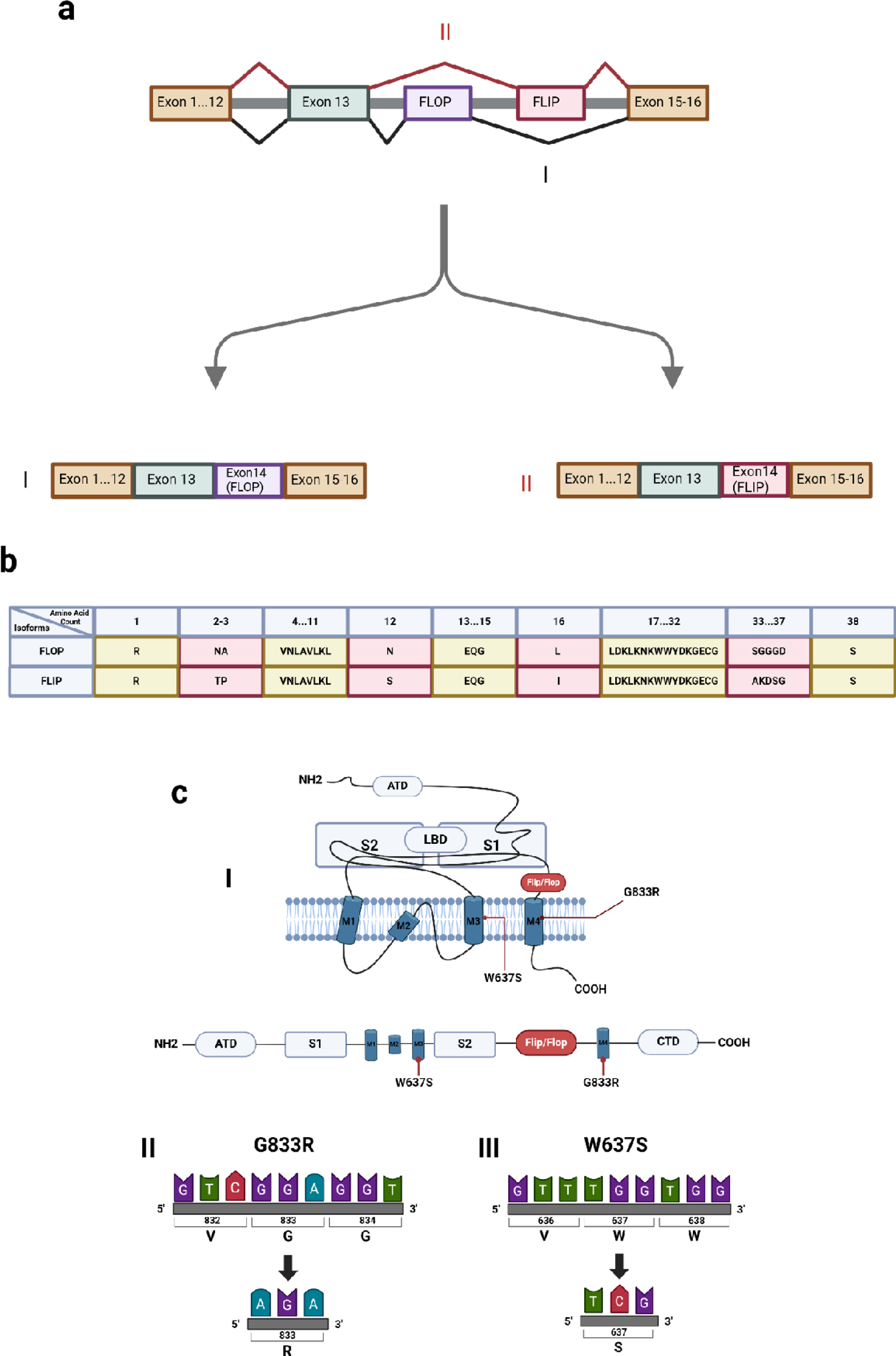
**a.** Illustration depicting exons spliced in two different ways through alternative splicing. **I)** The pathway causing the FLOP isoform is shown in a black line. When the FLOP isoform is formed, FLIP is skipped during splicing and binds to exon 15. **II)** The pathway causing the FLIP isoform is shown in the red line. In contrast to the FLOP isoform, in the FLIP isoform, the FLOP is skipped during splicing and binds to exon 15. **b.** Differences between the amino acids of the FLIP-FLOP isoforms of GRIA3 are illustrated. Amino acid groups that are the same are highlighted in yellow boxes, while amino acid groups that differ are highlighted in pink boxes. **c.** Modeling of variants **I)** The G833R substitution is located in transmembrane domain-4 (M4). The W637S substitution is seen in the transmembrane domain-3 (M3). Therefore, they are not affected by changes in FLIP-FLOP isoforms. **II)** A hemizygous change from G to A (c.2755G>A) is observed in the 15th exon of the GRIA3 gene located on the X chromosome. Thus, it results in the gly833arg (p.G833R) in amino acids. **III)** A hemizygous G to C (c.1910G>C) change is observed in the 12th exon of the GRIA3 gene located on the X chromosome. Thus, it results in trp637ser (p.W637S) in amino acids. (Figures are created in biorender.com)

Four missense variations were found in the functional domains of iGluR3 in experiments of Wu et al.: G833R, M706T, R631S, and R450Q. We are considering the G833R variant, which has the most effect. Studies on G833R expression in HEK293T (Human Embryonic Kidney) cells revealed that misfolding of the protein led to a 78% decrease in iGluR3. This variant is located in a critical region for iGlur3, in the transmembrane domain 4 (**Figure 1c**), and patients with this variation exhibit autistic characteristics, seizures, myoclonic convulsions, macrocephaly, and moderate mental retardation (Wu et al., 2007). We worked on both the G833R variant and the W637S variant. This variant is included in our study for the first time in the literature. The variant was first seen in a 2-year-old male patient. The patient’s family applied to the hospital with findings of developmental delay and muscle weakness, and the variant was detected through peripheral blood sample analysis. It has been classified as highly likely pathogenic by the genetic diagnosis center.

### iGLUR3 Expression Profile on Cell lines

To detect and indicate the presence and expression of iGLUR3 in cells and to select the appropriate cell line for future experiments, an iGLUR3 expression profile was generated. To compare the differences in iGLUR3 expression between neural and non-neural cell lines, a total of 7 cell lines, both neural and non-neural, were selected: HEK293T, RAJI, JURKAT, A549, A172, SHSY5Y, and U87. The antibody staining method was used to detect iGLUR3 expression in the cells. An antibody conjugated with Phycoerythrin (PE) and monoclonal was preferred, and flow cytometry analysis was performed using a Beckman Coulter CytoFlex instrument after staining the cells. During the flow cytometry measurement, the same cells were analyzed without staining as a control, in addition to the stained cells (Figure 2a). In the column chart depicting iGLUR3 expressions in the stained cells, it is evident that the iGLUR3 Mean Fluorescence Intensity (MFI) values of A172 cells are much higher than those of other cells (Figure 2b). However, it was observed that U87 and SHSY5Y cells exhibited lower iGLUR3 MFI values compared to the other cells. Our RT-qPCR analysis revealed significantly lower GRIA3 expression in SH-SY5Y and A172 cell lines compared to the control gene ESD exon 1. Specifically, SH-SY5Y cells showed minimal GRIA3 expression across exons 1, 3-4, 7-8, and 12-13, while A172 cells demonstrated low expression levels across the same exons. In contrast, U87-MG cells showed higher GRIA3 expression, particularly for exons 3-4 and 12-13, whereas no expression was detected in HEK293T cells (Figure 2c). Comparing our results with data from the Human Protein Atlas, we found consistency: SH-SY5Y and U87-MG cells generally exhibit low GRIA3 expression, whereas A172 and HEK293T cells show very low or no expression of GRIA3, consistent with our findings.

**Figure 2:**
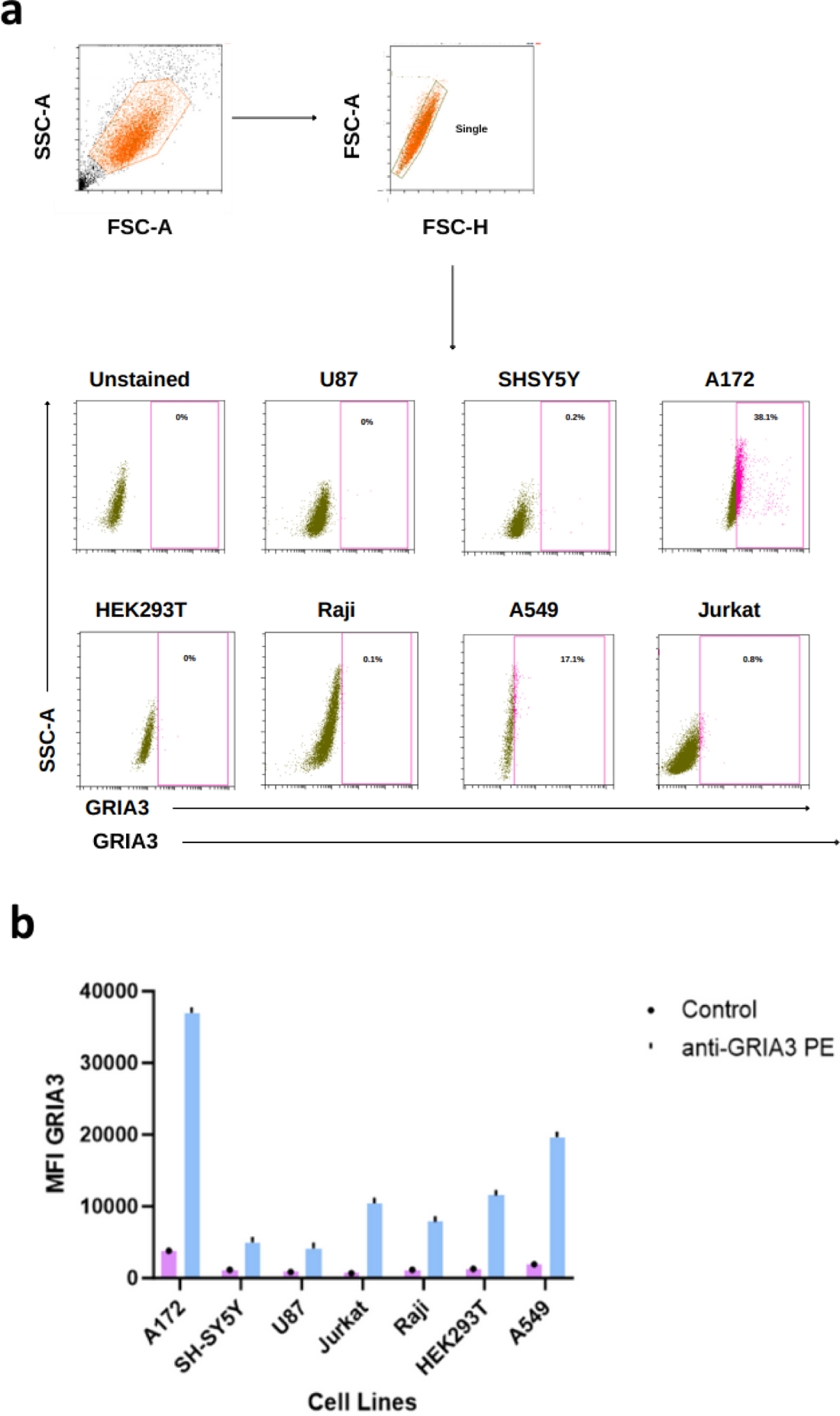
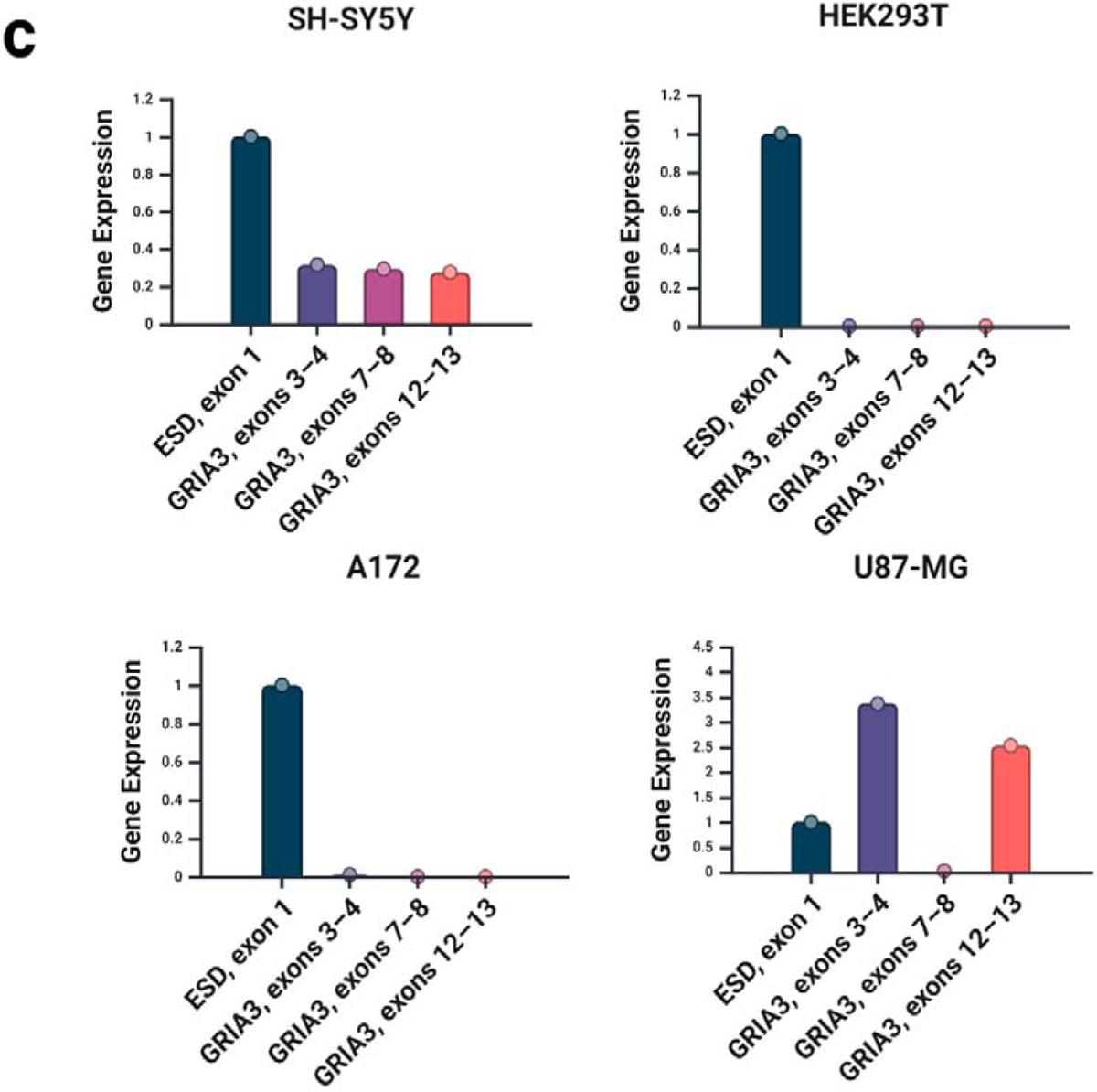
**a.** Demonstration of iGLUR3 expression obtained as a result of iGLUR3 antibody staining on flow cytometry. **b.** Bar graph showing GRIA3 MFI values obtained by flow cytometry analysis as a result of iGLUR3 antibody staining. **c.** The bar graph illustrates the gene expression levels of GRIA3 in non-transduced SH-SY5Y, HEK293T, A172, and U87-MG cells, using exon-specific primers and normalized to the ESD exon 1 expression.

### Production of FLIP, FLOP, G833R, and W637S Lentiviruses

After transfection with the isolated plasmids, host HEK293T cells were examined with the help of a fluorescent microscope under a GFP fluorescent filter for quality control of the success of the transfection. Since the cells carried the pHIV-EGFP tag cloned in the plasmid system, GFP fluorescence was observed as a result of transfection. This shows us that lentivirus production was successful (Figure 3a). To analyze the effectiveness of the lentivirus, a titration setup was established with decreasing concentrations. Three days after the setup was established, these Jurkat cells, to which lentivirus was applied with different concentrations, were analyzed by flow cytometry. Flow cytometry analysis was performed to examine transfection success and evaluate GFP expression. Compared to control JURKAT cells without lentivirus, in the presence of 10 µl lentivirus, JURKAT cells showed 93.04% GFP expression with FLIP lentivirus, 87.85% GFP expression with FLOP lentivirus, 48.52% GFP expression with G833R lentivirus, and 54.19% GFP expression with W637S lentivirus (Figure 3b). As the concentration of the lentivirus to which Jurkat cells are exposed decreases, the percentages of the cell population expressing GFP also decrease, confirming the success of lentivirus production (Figure 3c).

**Figure 3:**
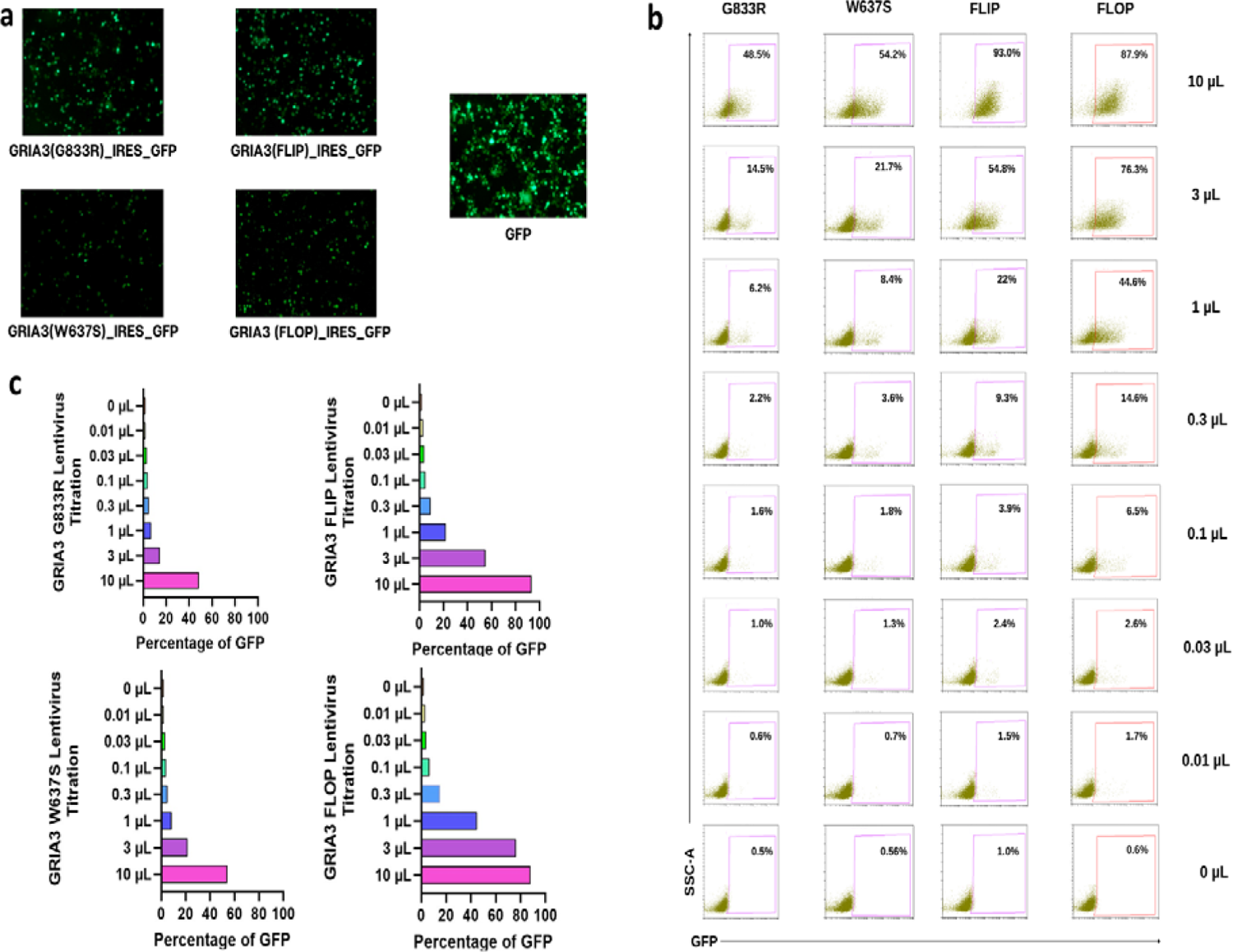
**a.** Images of HEK293T cells under a fluorescence microscope 3 days after transfection. **b.** Flow cytometry titration analysis three days after application of lentiviruses to Jurkat cells. **c.** Bar graph showing the percentages of GFP expression after application of different concentrations of lentiviruses on Jurkat cells.

### iGLUR3 Expression Profile on Wild-type Transduced Cells

Four gene sequences were designed: FLIP, FLOP isoforms, and G833R, W637S variants. These sequences were then cloned into the pHIV-EGFP plasmid after synthesis by genscript. To confirm that gene expression was absorbed into the cells, plasmids were preferred as pHIV-EGFP labeled. Thus, it could be confirmed that the cells receiving GFP (Green Fluorescent Protein) fluorescence were successfully transduced. To observe how transduction with wild-type changes the expression levels in the cells, the produced lentiviruses were first applied to JURKAT, U87, and A172 cells. Although both FLIP and FLOP isoforms are wild-type, we observed that the FLOP isoform was given priority as wild-type in experiments conducted in the literature. Therefore, to control the consistency of our experiments, priority was given to the FLOP isoform as wild-type and the first transductions were performed with the FLOP isoform. In addition, the cells were transduced with GFP, and a set of control groups were prepared. Transduced cells were subjected to antibody staining as before and flow cytometry analysis was performed. After FLOP and control GFP transduction in the cells, the cells were stained with iGLUR3 antibody processed with the GFP channel and analyzed with a flow cytometer system to demonstrate both iGLUR3 expression and GFP fluorescence. In the GRIA3-GFP quadrant graph, the percentages of cells showing GFP (+) GRIA3 (-), GFP (+) GRIA3 (+), GFP (-) GRIA3 (-), GFP (-) GRIA3 (+) characteristics were determined. Accordingly, the GFP (+) GRIA3 (+) percentage of FLOP transduced cells was observed as 5.95% in U87 cells, 19.97% in A172 cells, and 35.93% in JURKAT cells (Figure 4a). All of the staining we have done so far has shown iGLUR3 expression in the cell membrane. AMPA receptors, including iGLUR3, can be found inside the cell and are released into the cell membrane in the early LTP phase. For this reason, intracellular staining of the cells was performed with the Fixation/permeabilization procedure to demonstrate the expression of iGLUR3 within the cell. To do this, we started by staining JURKAT cells. When intracellular staining was performed on Jurkat cells, a GFP (+) GRIA3 (+) population was observed. This confirmed iGLUR3 expression inside Jurkat cells (Figure 4b). Since the cell populations were heterogeneous in terms of GFP and iGLUR3 expressions, single-cell sorting sets were established from transduced A172 FLOP, U87 GFP, U87 FLOP, U87 FLIP, U87 G833R, U87 W637S cell groups to select only GFP (+) GRIA3 (+) cells. Thus, cells with GFP expression levels above 95% were selected. As a result of the sorting process, GFP expression in each clone was analyzed by flow cytometry (Figure 4c). Following the cell sorting process, RT-qPCR was performed on these cells using specific primers. Among the transduced variants, U87 G833R showed the highest GRIA3 expression, followed by A172 FLOP, U87 FLOP, and U87 FLIP (Figure 4d). This significant increase in expression highlights the effectiveness of our lentiviral transduction in boosting GRIA3 expression. These consistent patterns validate the accuracy of our experimental approach. The increased expression in transduced cells indicates potential applications for these GRIA3 variants in functional studies, suggesting further research is warranted (Human Protein Atlas, 2024).

**Figure 4:**
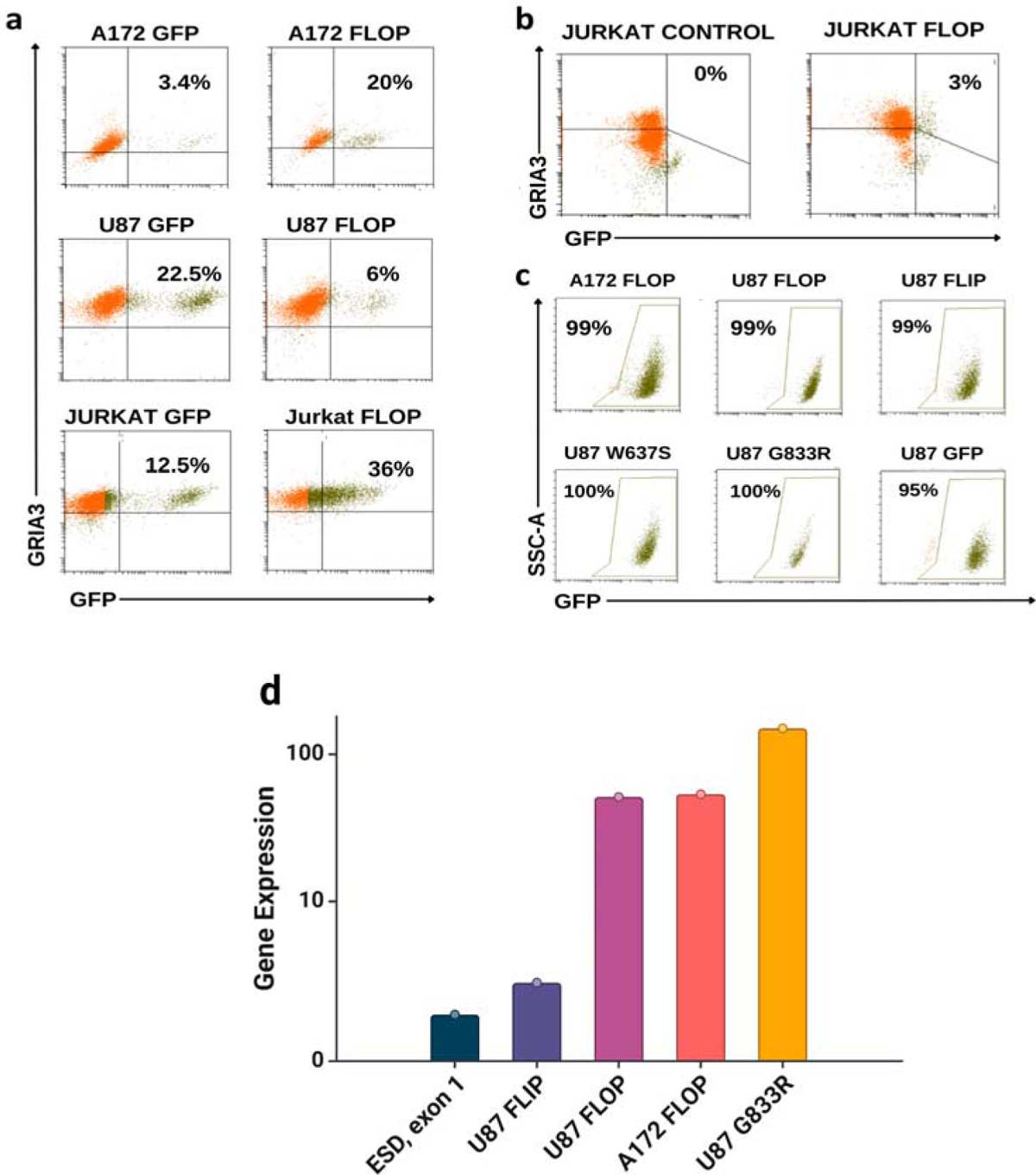
**a.** A quadrant graph showing the results of flow cytometry analysis with GRIA3 antibody staining and GFP channel processing in U87, A172, and JURKAT cells transduced with FLOP and GFP. **b.** A quadrant plot showing intracellular antibody staining of JURKAT cells transduced with the FLOP isoform. **c.** Demonstration of GFP expression in cell groups using single-cell sorting. **d.** Enhanced GRIA3 expression in transduced U87 (FLIP, FLOP, G833R) and A172 (FLOP) cells, highlighting the effectiveness of lentiviral transduction, with the G833R variant in U87 cells showing the highest expression.

## DISCUSSION

The G833R variant, mentioned in the study by Wu et al., and the W637S variant, included in our study for the first time, are just two examples of pathogenic variations. Some variants have not yet been researched. In addition, although not taken into account in the previous disease-related research, iGLUR3 has two isoforms, FLIP and FLOP, each causing changes in channel flow (Pei et al., 2007). Therefore, both of these isoforms were considered in our experiments aiming to detail the effects of isoforms.

The primary focus was on creating an iGLUR3 expression profile library by performing antibody staining on multiple cell lines. In this study, it was observed that iGLUR3, although primarily functioning in neurons, is also expressed in non-neural cells, possibly even more so than in neural cells. This made us consider that this illness might encompass more than just a mental retardation. The cells express iGLUR3, albeit at varying levels. To show this more clearly, when examining Mean Fluorescence Intensity (MFI) values, it was noted that the highest level of iGLUR3 expression was observed in the A172 cells among seven cell lines. Given that A172 is a neural cell line, this expression was anticipated. However, what was noteworthy is that certain non-neural cells exhibited higher expression levels than neural cells. These expression disparities may have implications for signaling pathways. It is aimed to perform antibody staining on additional cell lines to further expand the iGLUR3 expression profile library.

In our study, we observed varying levels of iGLUR3 protein expression in different cell lines through flow cytometry analysis. However, upon conducting RT-qPCR, we did not detect iGLUR3 expression in wild-type A172 cells. This discrepancy suggests that protein expression levels observed solely through flow cytometry antibody staining may not always correlate with mRNA expression levels, highlighting the need for caution in interpreting protein expression data from flow cytometry alone. It is possible that even low levels of mRNA expression could lead to detectable protein levels. Further investigations are warranted to elucidate the discrepancies between mRNA and protein expression levels.

AMPA receptors are found both in the cell membrane and inside the cell, and they are transported to the cell membrane with the assistance of kinases during the early phase of LTP, or they can be produced more during the late phase of LTP (Kandel et al., 2013).

The intracellular expression of AMPA receptors, including iGLUR3, raised questions. Therefore, this time, we focused on analyzing iGLUR3 expression within the cell. It was observed that the GFP (+) GRIA3 (+) population in the quadrant graph created during flow cytometry analysis correlated with the expression of iGLUR3 in JURKAT cells.

Thanks to the clones obtained from successfully transduced cells, we have created the first in vitro transgenic cell models, which will pave the way for genetic treatments of this disease, drug screening, and understanding its molecular mechanisms. To comprehend the mechanisms associated with mutations in GLUR3, mutant GLUR3 transgenic cell models were established. This allowed us to develop cell platforms suitable for future patch clamp analyses and neural cell modeling studies. Additionally, these experiments provided an opportunity to observe the effects of drugs on GLUR3 and genetic treatments on the transgenic cells created. These cells will serve as the foundation for all in vitro studies. In the subsequent study resulting from this work, selected genetic designs can be utilized in vivo and gene therapies.

We will report the differences between iGLUR3 expressions of variants and wild-type isoforms by performing antibody staining on these cells. Additionally, by conducting patch-clamp analysis on these cells, we will observe how wild-type isoforms and variants affect channel flow. Furthermore, signaling in non-neural cells can be observed to understand the function of iGLUR3 in non-neural cells via patch-clamp analysis. Questions such as whether iGLUR3 is active in non-neural cells and why iGLUR3 is expressed in non-neural cells are among those that need to be addressed in future studies. Among our plans, we aim to expand the iGLUR3 expression profile library by conducting antibody staining in more cells. We plan to delete iGLUR3 expression from cell lines using CRISPR-Cas9, analyze knockout cells, and restore normal iGLUR3 expression levels using CRISPR prime editing. This approach will facilitate the treatment of this disease.

## References

Abuin, Liliane, Benoîte Bargeton, Maximilian H. Ulbrich, Ehud Y. Isacoff, Stephan Kellenberger, ve Richard Benton. “Functional architecture of olfactory ionotropic glutamate receptors”. Neuron 69, sy 1 (13 Ocak 2011): 44–60. 10.1016/J.NEURON.2010.11.042.

Alt, Andrew, Eric S. Nisenbaum, David Bleakman, ve Jeffrey M. Witkin. “A role for AMPA receptors in mood disorders”. Biochemical Pharmacology 71, sy 9 (28 Nisan 2006): 1273–88. 10.1016/J.BCP.2005.12.022.

Authenticated HEK293T Cell Line Sigma Aldrich. Erişim 09 Ocak 2024. https://www.sigmaaldrich.com/TR/en/product/sigma/cb_12022001.

Blanke, Marie L., ve Antonius M.J. VanDongen. “Activation Mechanisms of the NMDA Receptor”. Biology of the NMDA Receptor, 01 Ocak 2009, 283–312. 10.1201/9781420044157.ch13.

Bonnet, C., B. Leheup, M. Béri, C. Philippe, M. J. Grégoire, ve P. Jonveaux. “Aberrant GRIA3 transcripts with multi-exon duplications in a family with X-linked mental retardation”. American Journal of Medical Genetics Part A 149A, sy 6 (01 Haziran 2009): 1280–89. 10.1002/AJMG.A.32858.

Chen, Xin, Lina Kozhaya, Cihan Tastan, Lindsey Placek, Mikail Dogan, Meghan Horne, Rebecca Abblett, vd. “Functional Interrogation of Primary Human T Cells via CRISPR Genetic Editing”. The Journal of Immunology 201, sy 5 (01 Eylül 2018): 1586–98. 10.4049/JIMMUNOL.1701616.

Chiyonobu, Tomohiro, Shin Hayashi, Kazuhiro Kobayashi, Masafumi Morimoto, Yuri Miyanomae, Akira Nishimura, Akemi Nishimoto, vd. “Partial tandem duplication of GRIA3 in a male with mental retardation”. American Journal of Medical Genetics Part A 143A, sy 13 (01 Temmuz 2007): 1448–55. 10.1002/AJMG.A.31798.

Felizola, Saulo J.A., Yasuhiro Nakamura, Fumitoshi Satoh, Ryo Morimoto, Kumi Kikuchi, Tomohiro Nakamura, Atsushi Hozawa, vd. “Glutamate receptors and the regulation of steroidogenesis in the human adrenal gland: The metabotropic pathway”. Molecular and Cellular Endocrinology 382, sy 1 (25 Ocak 2014): 170–77. 10.1016/J.MCE.2013.09.025.

Gándara, Carolina, Valerie Affleck, ve Elizabeth Ann Stoll. “Manufacture of Third-Generation Lentivirus for Preclinical Use, with Process Development Considerations for Translation to Good Manufacturing Practice”. Human gene therapy methods 29, sy 1 (01 Şubat 2018): 1–15. 10.1089/HGTB.2017.098.

Gonzalez, Jennifer, Vasanthi Jayaraman, Alexander S Maltsev, Robert E Oswald, Michael K Fenwick, Kinning Poon, ve Linda M Nowak. “2529-Pos Board B499”, 2009.

Guilmatre, A., Dubourg, C., Mosca, A. L., Legallic, S., Goldenberg, A., Drouin-Garraud, V., Layet, V., Rosier, A., Briault, S., Bonnet-Brilhault, F., Laumonnier, F., Odent, S., Le Vacon, G., Joly-Helas, G., David, V., Bendavid, C., Pinoit, J. M., Henry, C., Impallomeni, C., Germano, E., … Campion, D. (2009). Recurrent rearrangements in synaptic and neurodevelopmental genes and shared biologic pathways in schizophrenia, autism, and mental retardation. Archives of general psychiatry, 66(9), 947–956. 10.1001/archgenpsychiatry.2009.80

Guilmatre, Audrey, Christèle Dubourg, Anne Laure Mosca, Solenn Legallic, Alice Goldenberg, Valérie Drouin-Garraud, Valérie Layet, vd. “Recurrent rearrangements in synaptic and neurodevelopmental genes and shared biologic pathways in schizophrenia, autism, and mental retardation”. Archives of General Psychiatry 66, sy 9 (Eylül 2009): 947. 10.1001/ARCHGENPSYCHIATRY.2009.80.

Hamanaka, Kohei, Keita Miyoshi, Jia Hui Sun, Keisuke Hamada, Takao Komatsubara, Ken Saida, Naomi Tsuchida, vd. “Amelioration of a neurodevelopmental disorder by carbamazepine in a case having a gain-of-function GRIA3 variant”. Human Genetics 141, sy 2 (01 Şubat 2022): 283–93. 10.1007/S00439-021-02416-7/FIGURES/4.

Hansen, Kasper B., Feng Yi, Riley E. Perszyk, Hiro Furukawa, Lonnie P. Wollmuth, Alasdair J. Gibb, ve Stephen F. Traynelis. “Structure, function, and allosteric modulation of NMDA receptors”. Journal of General Physiology 150, sy 8 (06 Ağustos 2018): 1081–1105. 10.1085/JGP.201812032.

Henley, Jeremy M., ve Kevin A. Wilkinson. “AMPA receptor trafficking and the mechanisms underlying synaptic plasticity and cognitive aging”. Dialogues in Clinical Neuroscience 15, sy 1 (Mart 2013): 11–27. 10.31887/DCNS.2013.15.1/JHENLEY.

Human Protein Atlas. (2024). Expression of GRIA3 in cell lines. Retrieved from https://www.proteinatlas.org.

Karakas, Erkan, ve Hiro Furukawa. “Crystal structure of a heterotetrameric NMDA receptor ion channel”. Science 344, sy 6187 (30 Mayıs 2014): 992–97. 10.1126/SCIENCE.1251915/SUPPL_FILE/1251915.KARAKAS.SM.PDF.

Kato, Akihiko S., Martin B. Gill, Hong Yu, Eric S. Nisenbaum, ve David S. Bredt. “TARPs differentially decorate AMPA receptors to specify neuropharmacology”. Trends in Neurosciences 33, sy 5 (01 Mayıs 2010): 241–48. 10.1016/J.TINS.2010.02.004.

Kew, James N.C., ve John A. Kemp. “Ionotropic and metabotropic glutamate receptor structure and pharmacology”. Psychopharmacology 179, sy 1 (25 Nisan 2005): 4–29. 10.1007/S00213-0052200-Z/FIGURES/13.

Kleiveland, Charlotte, ve Charlotte Kleiveland. “Peripheral Blood Mononuclear Cells”. The Impact of Food Bioactives on Health: In Vitro and Ex Vivo Models, 01 Ocak 2015, 161–67. 10.1007/978-3-319-16104-4_15.

Kordi-Tamandani, Dor Mohammad, Nahid Dahmardeh, ve Adam Torkamanzehi. “Evaluation of hypermethylation and expression pattern of GMR2, GMR5, GMR8, and GRIA3 in patients with schizophrenia”. Gene 515, sy 1 (15 Şubat 2013): 163–66. 10.1016/J.GENE.2012.10.075.

Kovalevich, Jane, ve Dianne Langford. “Considerations for the Use of SH-SY5Y Neuroblastoma Cells in Neurobiology”. Methods in molecular biology (Clifton, N.J.) 1078 (2013): 9. 10.1007/978-1-62703-640-5_2.

Kukushkin, Nikolay Vadimovich, ve Thomas James Carew. “Memory Takes Time”. Neuron 95, sy 2 (19 Temmuz 2017): 259–79. 10.1016/J.NEURON.2017.05.029.

Lau, Anthony, ve Michael Tymianski. “Glutamate receptors, neurotoxicity and neurodegeneration”. Pflugers Archiv European Journal of Physiology 460, sy 2 (14 Temmuz 2010): 525–42. 10.1007/S00424-010-0809-1/METRICS.

Lü Scher, Christian, Robert C Malenka, Morgan Sheng, Bernardo Sabatini, ve Thomas C Sü. “NMDA Receptor-Dependent Long-Term Potentiation and Long-Term Depression (LTP/LTD)”. Cold Spring Harbor Perspectives in Biology 4, sy 6 (01 Haziran 2012): a005710. 10.1101/CSHPERSPECT.A005710.

M. Karadağ. (2017). Mental Retardasyonu Olan Çocuk ve Ergenlerin Tedavisinde Kullanılan Farmakolojik Ajanlar. Çocuk ve Gençlik Ruh Sağlığı Dergisi, 3(24), Article 70685.

Martinez-Esteve Melnikova, Anastasia, Jordi Pijuan, Javier Aparicio, Alia Ramírez, Anna Altisent-Huguet, Alba Vilanova-Adell, Alexis Arzimanoglou, vd. “The p.Glu787Lys variant in the GRIA3 gene causes developmental and epileptic encephalopathy mimicking structural epilepsy in a female patient”. European Journal of Medical Genetics 65, sy 3 (01 Mart 2022): 104442. 10.1016/J.EJMG.2022.104442.

McManus, C. Joel, ve Brenton R. Graveley. “RNA structure and the mechanisms of alternative splicing”. Current opinion in genetics & development 21, sy 4 (Ağustos 2011): 373–79. 10.1016/J.GDE.2011.04.001.

Merten, Otto Wilhelm, Matthias Hebben, ve Chiara Bovolenta. “Production of lentiviral vectors”. Molecular Therapy Methods & Clinical Development 3 (01 Ocak 2016): 16017. 10.1038/MTM.2016.17.

Newcomer, John W., Nuri B. Farber, ve John W. Olney. “NMDA receptor function, memory, and brain aging”. Dialogues in Clinical Neuroscience 2, sy 3 (30 Eylül 2000): 219. 10.31887/DCNS.2000.2.3/JNEWCOMER.

Okano, S., Makita, Y., Miyamoto, A., Taketazu, G., Kimura, K., Fukuda, I., Tanaka, H., Yanagi, K., & Kaname, T. (2023). GRIA3 p.Met661Thr variant in a female with developmental epileptic encephalopathy. Human genome variation, 10(1), 4. 10.1038/s41439-023-00232-1

Paoletti, Pierre. “Molecular basis of NMDA receptor functional diversity”. The European journal of neuroscience 33, sy 8 (2011): 1351–65. 10.1111/J.1460-9568.2011.07628.X.

Pei, Weimin, Zhen Huang, Congzhou Wang, Yan Han, Seon Park Jae, ve Li Niu. “Flip and flop: A molecular determinant for AMPA receptor channel opening”. Biochemistry 48, sy 17 (05 Mayıs 2009): 3767–77. 10.1021/BI8015907/ASSET/IMAGES/MEDIUM/BI-2008-015907_0007.GIF.

Philips, A. K., Sirén, A., Avela, K., Somer, M., Peippo, M., Ahvenainen, M., Doagu, F., Arvio, M., Kääriäinen, H., Van Esch, H., Froyen, G., Haas, S. A., Hu, H., Kalscheuer, V. M., & Järvelä, I. (2014). X-exome sequencing in Finnish families with intellectual disability--four novel mutations and two novel syndromic phenotypes. Orphanet journal of rare diseases, 9, 49. 10.1186/1750-1172-9-49

Philips, Anju K., Auli Sirén, Kristiina Avela, Mirja Somer, Maarit Peippo, Minna Ahvenainen, Fatma Doagu, vd. “X-exome sequencing in Finnish families with Intellectual Disability - four novel mutations and two novel syndromic phenotypes”. Orphanet Journal of Rare Diseases 9, sy 1 (11 Nisan 2014): 49. 10.1186/1750-1172-9-49.

Piard, Juliette, Matthieu Béreau, Wenshu XiangWei, Thomas Wirth, Daniel Amsallem, Lauren Buisson, Philippe Richard, vd. “The GRIA3 c.2477G > A Variant Causes an Exaggerated Startle Reflex, Chorea, and Multifocal Myoclonus”. Movement disordersC: official journal of the Movement Disorder Society 35, sy 7 (01 Temmuz 2020): 1224. 10.1002/MDS.28058.

Pourahmad, Jalal, ve Ahmad Salimi. “Isolated Human Peripheral Blood Mononuclear Cell (PBMC), a Cost Effective Tool for Predicting Immunosuppressive Effects of Drugs and Xenobiotics”. Iranian Journal of Pharmaceutical ResearchC: IJPR 14, sy 4 (01 Eylül 2015): 979.

Purves, Dale, George J Augustine, David Fitzpatrick, Lawrence C Katz, Anthony-Samuel LaMantia, James O McNamara, ve S Mark Williams. “Glutamate Receptors”, 2001. https://www.ncbi.nlm.nih.gov/books/NBK10802/.

Riedel, Gernot, Bettina Platt, ve Jacques Micheau. “Glutamate receptor function in learning and memory”. Behavioural Brain Research 140, sy 1-2 (18 Mart 2003): 1–47. 10.1016/S0166-4328(02)00272-3.

Song, Insuk, ve Richard L. Huganir. “Regulation of AMPA receptors during synaptic plasticity”. Trends in Neurosciences 25, sy 11 (01 Kasım 2002): 578–88. 10.1016/S0166-2236(02)02270-1.

Sun, Jia Hui, Jiang Chen, Fernando Eduardo Ayala Valenzuela, Carolyn Brown, Diane Masser-Frye, Marilyn Jones, Leslie Patron Romero, vd. “X-linked neonatal-onset epileptic encephalopathy associated with a gain-of-function variant p.R660T in GRIA3”. PLoS genetics 17, sy 6 (23 Haziran 2021). 10.1371/JOURNAL.PGEN.1009608.

Tajima, Nami, Erkan Karakas, Timothy Grant, Noriko Simorowski, Ruben Diaz-Avalos, Nikolaus Grigorieff, ve Hiro Furukawa. “Activation of NMDA receptors and the mechanism of inhibition by ifenprodil”. Nature 2016 534:7605 534, sy 7605 (02 Mayıs 2016): 63-68. 10.1038/nature17679.

Tastan, Cihan, Ece Karhan, Wei Zhou, Elizabeth Fleming, Anita Y. Voigt, Xudong Yao, Lei Wang, vd. “Tuning of human MAIT cell activation by commensal bacteria species and MR1-dependent T-cell presentation”. Mucosal Immunology 11, sy 6 (01 Kasım 2018): 1591–1605. 10.1038/S41385-018-0072-X.

Taştan, Cihan, Derya Dilek Kançağı, Raife Dilek Turan, Bulut Yurtsever, Didem Çakırsoy, Selen Abanuz, Muhammet Yılancı, vd. “Preclinical Assessment of Efficacy and Safety Analysis of CAR-T Cells (ISIKOK-19) Targeting CD19-Expressing B-Cells for the First Turkish Academic Clinical Trial with Relapsed/Refractory ALL and NHL Patients”. Turkish journal of haematologyC: official journal of Turkish Society of Haematology 37, sy 4 (2020): 234–47. 10.4274/TJH.GALENOS.2020.2020.0070.

Traynelis, Stephen F., Lonnie P. Wollmuth, Chris J. McBain, Frank S. Menniti, Katie M. Vance, Kevin K. Ogden, Kasper B. Hansen, Hongjie Yuan, Scott J. Myers, ve Ray Dingledine. “Glutamate Receptor Ion Channels: Structure, Regulation, and Function”. Pharmacological Reviews 62, sy 3 (Eylül 2010): 405. 10.1124/PR.109.002451.

Vandenberghe, Wim, Roger A. Nicoll, ve David S. Bredt. “Stargazin is an AMPA receptor auxiliary subunit”. Proceedings of the National Academy of Sciences of the United States of America 102, sy 2 (11 Ocak 2005): 485–90. 10.1073/PNAS.0408269102/ASSET/2AE44ECD-07A6-4E7D-B641-1113B28518FF/ASSETS/GRAPHIC/ZPQ0510467660004.JPEG.

Wang, Y., Liu, J., Huang, B., Xu, Y., Li, J., Huang, L., Wang, X. (2015). Mechanism of alternative splicing and its regulation (Review). Biomedical Reports, 3, 152–158. 10.3892/br.2014.407

Whibley, Annabel C., Vincent Plagnol, Patrick S. Tarpey, Fatima Abidi, Tod Fullston, Maja K. Choma, Catherine A. Boucher, vd. “Fine-Scale Survey of X Chromosome Copy Number Variants and Indels Underlying Intellectual Disability”. American Journal of Human Genetics 87, sy 2 (08 Ağustos 2010): 173. 10.1016/J.AJHG.2010.06.017.

Wu, Ye, Amy C. Arai, Gavin Rumbaugh, Anand K. Srivastava, Gillian Turner, Takashi Hayashi, Erika Suzuki, vd. “Mutations in ionotropic AMPA receptor 3 alter channel properties and are associated with moderate cognitive impairment in humans”. Proceedings of the National Academy of Sciences of the United States of America 104, sy 46 (13 Kasım 2007): 18163–68. 10.1073/PNAS.0708699104.

Zhou, Qiang, ve Morgan Sheng. “NMDA receptors in nervous system diseases”. Neuropharmacology 74 (01 Kasım 2013): 69–75. 10.1016/J.NEUROPHARM.2013.03.030.

